# Proformer: a hybrid macaron transformer model predicts expression values from promoter sequences

**DOI:** 10.1101/2023.03.10.532129

**Authors:** Il-Youp Kwak, Byeong-Chan Kim, Juhyun Lee, Daniel J. Garry, Jianyi Zhang, Wuming Gong

**Author notes:** Co-corresponding authors: Wuming Gong, Ph.D., Lillehei Heart Institute, 2231 6^th^ St SE, University of Minnesota, Minneapolis, MN 55455, Daniel J. Garry, M.D., Ph.D., Lillehei Heart Institute, 2231 6^th^ St SE, University of Minnesota Minneapolis, MN 55455.

## Abstract

The breakthrough high-throughput measurement of the cis-regulatory activity of millions of randomly generated promoters provides an unprecedented opportunity to systematically decode the cis-regulatory logic that determines the expression values. We developed an end-to-end transformer encoder architecture named Proformer to predict the expression values from DNA sequences. Proformer used a Macaron-like Transformer encoder architecture, where two half-step feed forward (FFN) layers were placed at the beginning and the end of each encoder block, and a separable 1D convolution layer was inserted after the first FFN layer and in front of the multi-head attention layer. The sliding *k*-mers from one-hot encoded sequences were mapped onto a continuous embedding, combined with the learned positional embedding and strand embedding (forward strand vs. reverse complemented strand) as the sequence input. Moreover, Proformer introduced multiple expression heads with mask filling to prevent the transformer models from collapsing when training on relatively small amount of data. We empirically determined that this design had significantly better performance than the conventional design such as using the global pooling layer as the output layer for the regression task. These analyses support the notion that Proformer provides a novel method of learning and enhances our understanding of how cis-regulatory sequences determine the expression values.

## Introduction

Gene expression is a fundamental process and is essential for the coordinated function of all living organisms. Predicting the expression level of a gene based on its promoter or enhancer sequences is an important problem in molecular biology, with applications ranging from understanding the regulation of gene expression to engineering gene expression for biotechnological applications^1,2^. Recent progress and mechanistic insights have been obtained using large-scale and high-throughput massively parallel reporter assays (MPRAs), which enable the study of gene expression and regulatory elements in a high-throughput manner and the simultaneous testing of thousands to millions of enhancers or promoters in parallel^3–24^. MPRA protocols linked random or mutated sequences to unique barcodes, with each sequence-barcode pair represented in a different reporter assay vector. After delivery of the pooled vector library, barcode abundance could be subsequently quantified using next-generation sequencing (NGS) techniques^25^. MPARs enabled large scale studies of functional annotation of putative regulatory elements^3,26^, variant effect prediction^22,23,27,28^ and evolutionary reconstructions^25,29,30^. For example, STARR-seq (self-transcribing active regulatory region sequencing) was used to investigate the enhancer activities of tens of millions of independent fragments from the Drosophila genome^3^. Microarray-based or PCA-based (polymerase cycling assembly) synthesized DNA regulatory elements with unique sequence tags were used to evaluate hundreds of thousands of variants of mammalian promoters or enhancers^4–6^. Nguyen et al. systematically compared the promoter and enhancer potentials of many candidate sequences^10^. Using Gigantic Parallel Reporter Assay (GPRA), de Boer et al. measured the expression level associated with tens of millions of random promoter sequences and used these to learn cis-regulatory logic in the yeast grown in well-characterized carbon sources^14^.

Machine learning methods have been developed to identify complex relationships and patterns in large scale DNA sequences (including MPRA data) that may not be apparent through conventional statistical methods. For example, convolutional neural networks (CNN) and recurrent neural networks (RNN) were used to capture the local dependences in DNA sequences and/or genomic features and predict binding affinities^1,31,32^, chromatin features^33,34^, DNA methylation^35,36^, RBP (RNA-binding protein) binding^37–39^ and gene expression levels^40^. In contrast, Transformers are a type of neural network architecture that has gained popularity in recent years for their ability to process sequential data, such as text and speech, more efficiently and effectively than traditional RNNs and CNNs^41^. Transformers used an attention mechanism to selectively focus on different aspects of the input sequence, which allowed them to capture long-range dependencies more effectively than RNNs and CNNs that typically rely on fixed-length windows or sliding windows.

In this study, we developed an end-to-end transformer encoder architecture, Proformer, to predict the expression values from millions of DNA sequences. Our method introduces several innovative designs such as Macaron-like encoder structures, *k*-mer embedding, and multiple expression heads (MEH) to learn the relationships between a large number of sequences and expression values. Proformer ranked in the 3rd place in the final standing of the DREAM challenge: predicting gene expression using millions of random promoter sequences^42^. We believe that our model provides a novel method of learning and characterizing how cis-regulatory sequences determine the expression values. Codes pertaining to important analyses in this study are available from GitHub webpage: https://github.com/gongx030/dream_PGE.

## Results

### Proformer overview

Proformer used a Macaron-like Transformer encoder architecture to predict the expression values from promoter sequences (Figure 1)^43–45^. Compared with the regular Transformer encoder, the Macaron-like encoder has two half-step feed forward (FFN) layers at the beginning and the end of each Transformer encoder block, which can be mathematically interpreted as a numerical Ordinary Differential Equation (ODE) solver for a convection-diffusion equation in a multi-particle dynamic system^46,47^. Given the stochastic nature of the input sequences, we hypothesized that this design may better recover the associations between nucleotide pattern and the expression values. We added a separable 1D convolution layer in the Macaron encoder block following the first FFN layer and in front of the multi-head attention layer. This design has been used in other Transformer architectures such as Conformer^47^, and is shown to be critical for capturing the local signals.

**Figure 1.**
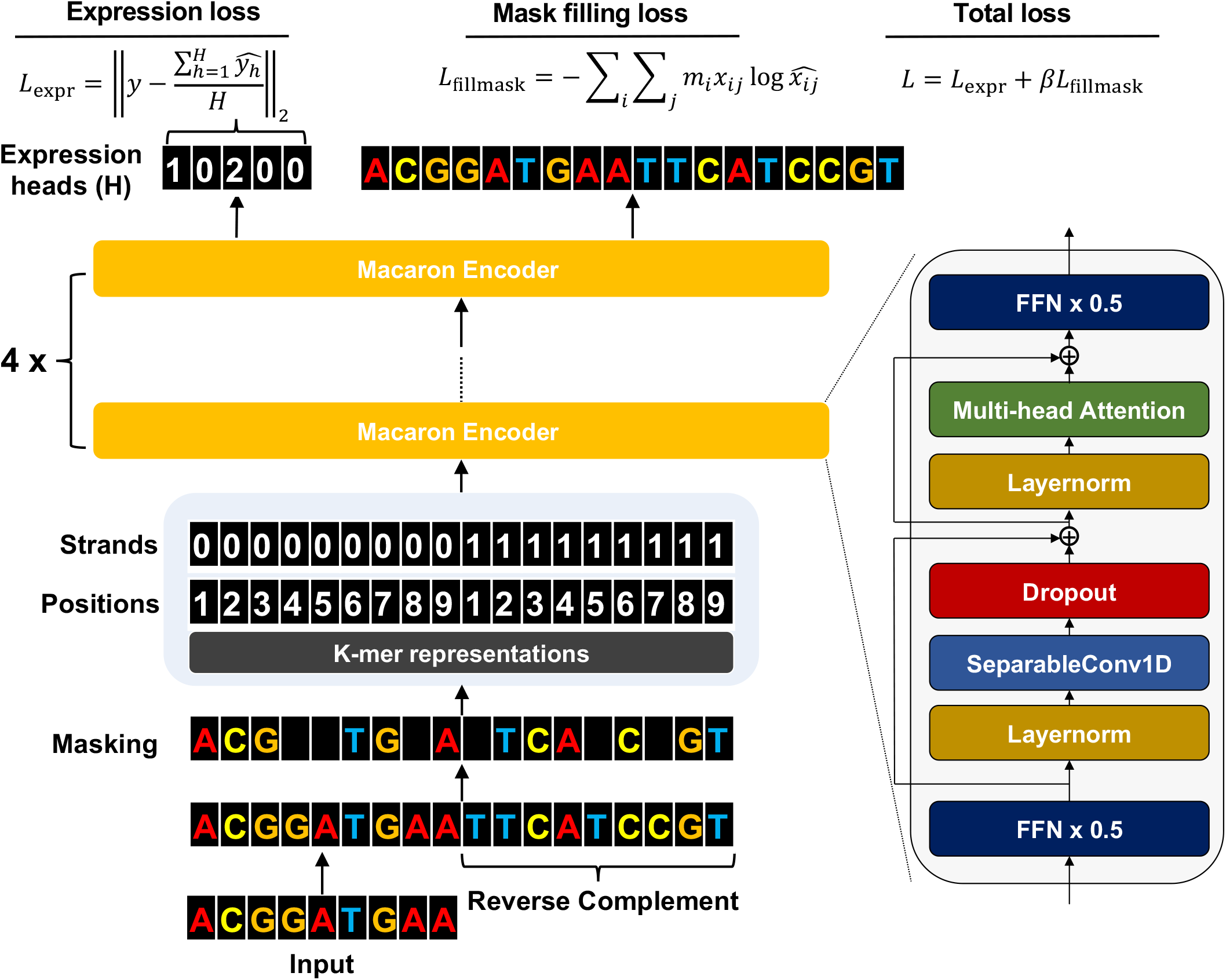
Proformer is a macaron-like transformer architecture that models the relationship between DNA sequences and expression values.

We extracted the sliding *k*-mers (*k*=10 in the final model) from one-hot encoded sequences and mapped them onto a continuous embedding platform. It has been previously shown that the *k*-mer embedding of nucleotide sequences had better performance than the convolution on tasks such as predicting transcription factor binding sites^48^. The *k*-mer embedding was then combined with the learned positional embedding and strand embedding (forward strand vs reverse complemented strand) as one part of the input to the Macaron encoder.

We added *H* positions (*H* = 32 in our final model) as the expression heads (Figure 1). Proformer predicted one expression value for each expression head and used the mean of the prediction of all positions as the final predicted expression value. The total training losses consisted of the mean squared error between predicted and observed expression values (*L_expr_*), and the reconstruction loss (*L_recon_*), where we randomly masked 5% of the nucleotides and had the model predict the masked nucleotides. In our final model, we set the weight for reconstruction loss *β* = 1.

The final Proformer model had approximately 47 million trainable parameters, implemented by TensorFlow 2 and trained on one machine with four A100 GPUs. We varied the learning rate over the course of training according to the formula used in the original Transformer paper^43^. Warmup steps of 12,500 and a batch size of 512 were used in the training. We used the Adam optimizer with *β*_1_ = 0.9, *β*_2_ = 0.98 and *ϵ* = 10”^9^ for these studies.

### MEH with mask filling has improved performance using large over-parameterized models

Global average pooling layer at the top of a neural network is commonly used for the regression and classification tasks^49^. However, we found that when applying the global average pooling layer at the top of a large transformer model, for example, with a dimension size of 256 and blocks size of 8, the whole model sometimes failed to converge on training on relatively small amount (~500k) of samples (Figure 2b). In order to address this issue, we proposed a new design, where the model predicted multiple expression values through multiple expression heads (MEH) and used the average of all predictions as the final predicted value (Figure 2a), while at the same time, the model also predicted the randomly masked DNA nucleotide. MEH with mask filling produced stable convergence when training the transformer model with the same size on ~500k samples (Figure 2b). In order to systematically compare the performance of two designs, we trained the models on 10% of the training sequence / expression value pairs then the performance was evaluated on 2% of the data as the Pearson’s R between observed and predicted expression values. For MEH with mask filling, we also examined the performance over a different number of heads (*H* = 1,8,16,32,64). Overall, we found that MEH with mask filing gave significantly better than global average pooling when using 8 or more heads (Mann-Whitney U test p-values = 0.0715, 0.0102, 0.0142, 0.0224 and 0.00605 for *H* = 1,8,16,32,64, respectively), and the best performance was achieved at a dimension size of 128 and macaron block size of 8 (Figure 2c). As the model size became larger and deeper, the global average pooling became difficult to converge, while in comparison, MEH with masking filing could still provide stable results.

**Figure 2.**
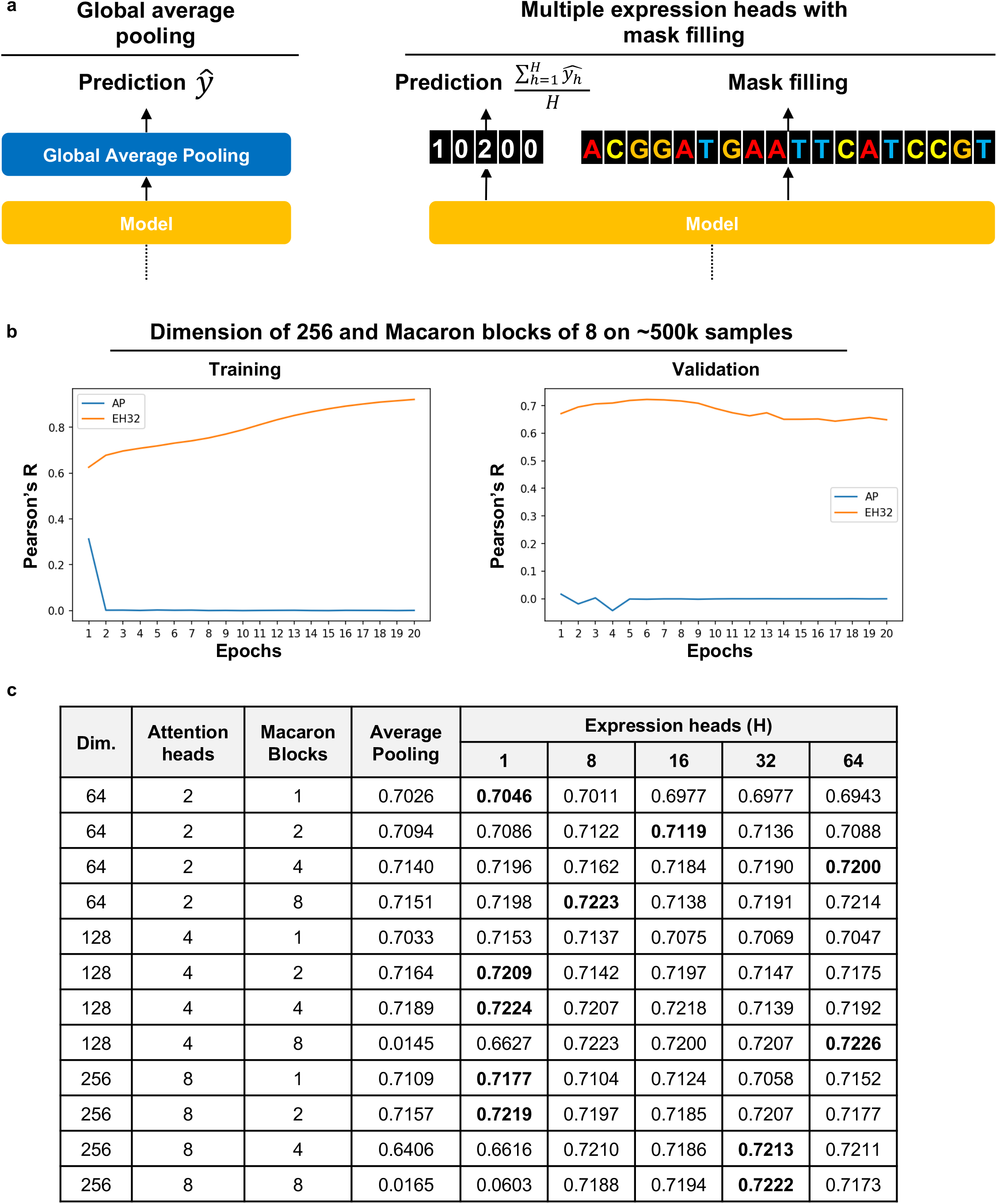
Multiple expression heads (MEH) with mask filling has better performance on large over-parameterized models. **(a)** Global average pooling layer and MEH with mask filling were used at the top the transformer blocks. **(b)** The training (left) and validation (left) performance of Proformer models using global average pooling (AP) or MEH with 32 heads (EH32) were compared. The performance was measured by the Pearson’s R between observed and predicted expression values. **(c)** Systematic evaluation of global average pooling and MEH with mask filing on different model specifications such as dimension heads (2, 4, and 8), macaron blocks (1, 2, 4, and 8), and number of expression heads (1, 8, 16, 32, 64) was performed. The best performance of each model specification was highlighted.

### MEH with mask filling has better performance for the prediction of chromatin accessibility from DNA sequences

To test whether our observations on these two head designs could apply to similar scenarios, we designed another task to use Proformer to predict ATAC-seq (Assay for Transposase-Accessible Chromatin with high-throughput sequencing) signals from DNA sequences. The ATAC-seq is a technique to measure the chromatin accessibility across the whole genome^50^. We sampled a total of 100k genomic sub-regions surrounding the ~80,000 summits of ATAC-seq data of GM12878^50^, while each genomic sub-region included 100 nucleotides. Different models were built to predict the mean ATAC-seq signal of the central 20 bp from 100 nt DNA sequences (Figure 3a). The global average pooling performed well when the model size was relatively small. As the model size became larger, we observed similar trends such that the global average pooling tended to fail on large over-parameterized models. The best performance was achieved by using MEH with mask filling with dimension size of 128 and block size of 4 (Figure 3b).

**Figure 3.**
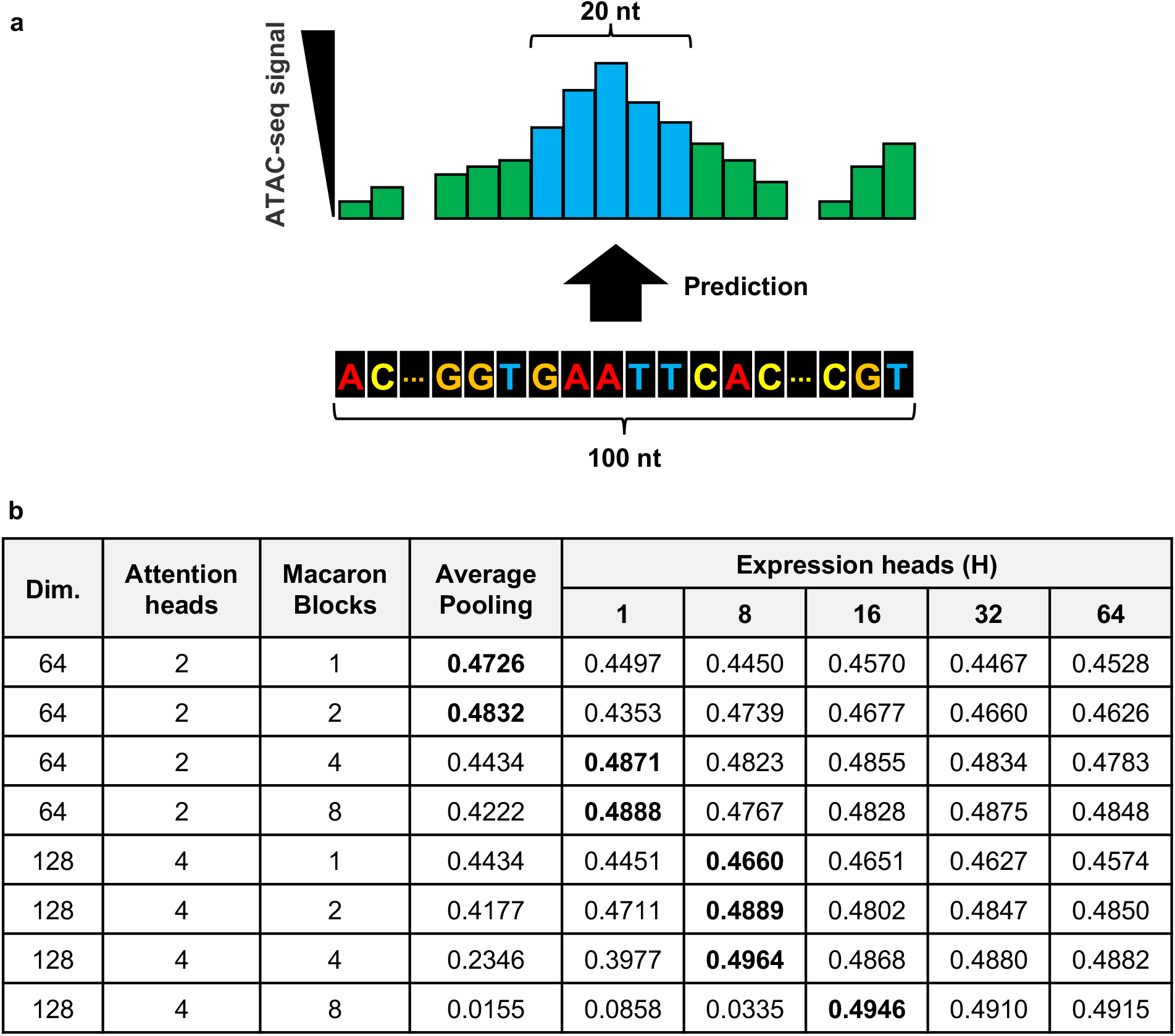
Multiple expression heads (MEH) with mask filling has better performance on predicting chromatin accessibility from DNA sequences. **(a)** The task of predicting mean ATAC-seq signal of the central 20 bp from 100 nt surrounding DNA sequences was examined. **(b)** Systematic evaluation of global average pooling and MEH with mask filing on different model specifications such as dimension heads (2 and 4), macaron blocks (1, 2, 4, and 8), and number of expression heads (1, 8, 16, 32, 64). The best performance of each model specification was highlighted.

### MEH with mask filling is critical for improving the prediction performance on hold-out validation data

We trained the final model for the DREAM challenge by using a dimension size of 512 and a block size of 4 on 95% of the data provided by the organizers and evaluated on the remaining 5%. The checkpoint after the 6th epoch was used where the validation Pearson’s R was maximized. As expected, MEH with mask filling produced improved Pearson’s R than global average pooling on the validation data (Figure 4a). The ablation study showed when using only one expression head (*H* = 1), the performance was similar to global average pooling. However, MEH with *H* = 32 showed improvement over hold-out validation data and produced the highest weighted scores. It is interesting that adding a GLU activation^51^ to expression heads produced even higher unweighted Pearson’s R and Spearman’s Rho on the hold-out validation data, while the weighted score became worse than global average pooling (Figure 4b). Future studies will explore different designs of the expression heads.

**Figure 4.**
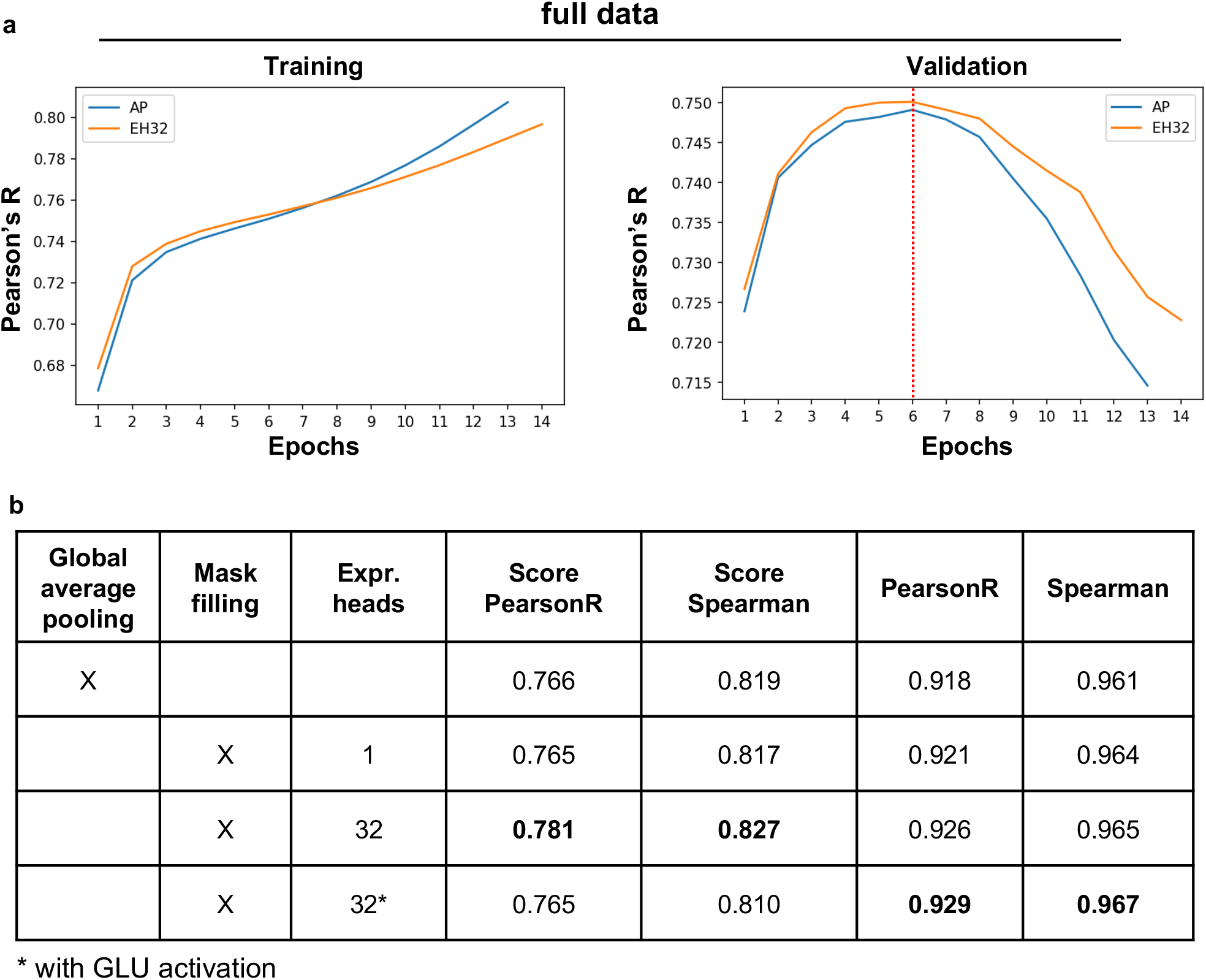
Multiple expression heads (MEH) with mask filling is critical for improving the prediction performance on hold-out validation data. **(a)** The training (left) and validation (left) performance of Proformer models on the full DREAM dataset using global average pooling (AP) or MEH with 32 heads (EH32). The performance was measured by the Pearson’s R between observed and predicted expression values. The checkpoint after the 6th epoch was used as the final model where the validation Pearson’s R was maximized (red dotted line). **(b)** The performance of Proformer model on the hold-out validation data. The performance is measured by weighted (score) or unweighted Pearson’s R and Spearman’s Rho between observed and predicted expression values.

## Discussion

Various machine learning techniques have been used to analyze and interpret the MPRA data and dissect the regulatory logics. Recently, over-parameterized deep networks or large models, with more parameters than the size of the training data, have dominated the performance in various machine learning areas^52^. The global average pooling layer was conventionally used to aggregate the information from multiple channels and to produce final predictions. However, we found that when training over-parameterized models on the regression tasks such as predicting expression values from DNA sequences, the global average pooling often led to a convergence issue, most likely due to the loss of information that accumulated during the training and caused the model to perform poorly or failed to converge. Here we presented a new architecture Proformer for prediction of expression values from DNA sequences. We introduced a new design named multiple expression heads (MEH) with mask filling to prevent the over-parameterized transformer models from collapsing when training on relatively small amount of data. Applying the Proformer model to predictexpression values and to predict chromatin accessibility from DNA sequences showed that MEH with masking filling produced significantly better performance and stable convergence compared to the commonly used global average pooling. Based on our studies, we propose that MEH with mask filling will be a useful design for similar regression tasks that took advantage of large over-parameterized models.

## Methods

### DREAM challenge dataset overview

Rafi et al. conducted a high-throughput experiment to measure the regulatory effect of millions of random DNA sequences. They cloned 80 bp random DNA sequences into a promoter-like context upstream of a yellow fluorescent protein (YFP), transformed the resulting library into yeast, and measured expression by fluorescent activated cell sorting^4,14,53^. The training dataset includes 6,739,258 random promoter sequences and their corresponding mean expression values^42^.

Rafi et al. also provided 71,103 sequences from several promoter sequence types as the hold-out “validation” dataset to evaluate the model performance in different ways. These validation datasets included predicting the expression changes resulting from single nucleotide variants (SNVs), perturbation of specific transcription factor (TF) binding sites, tiling of TF binding sites across background sequences, sequences with high- and low-expression levels, native yeast genomic sequences, random DNA sequences, and challenging sequences designed to maximize differences between a convolutional model and a biochemical model trained on the same data^42^.

### Sequence trimming and padding

We removed the leading 17 and trailing 13 nucleotides (nt) that were identical in both training and testing promoter sequences, since these nucleotides were not informative for the prediction of expression values and removal of the nucleotides would significantly reduce the training and inference time. The length of the resulting promoter sequences ranged from 48 to 112 nt for training data, while >99.97% training promoters were less than 100 nucleotides. To further reduce the computational overhead, we used 6,737,568 promoter sequences shorter than 100 nt (after trimming) in the model training. For promoters that were less than 100 nt, the left and right sides were padded with the letter N.

### Reverse complemented sequences

We empirically found that including the reverse complemented promoter sequences would significantly improve the performance. Thus, the reverse complemented sequences were concatenated with the original sequences (after trimming and padding) and used as the input for model training. Thus, the total length of the input sequences was 200 nt.

### Standardization of the expression values

The expression values were standardized to the mean of zero and standard deviation of one. Our experiments found that when the mean squared error loss was used, standardizing the expression values gave better model generalization performance (in terms of Pearson’s R and Spearman’s Rho) and faster convergence.

### ATAC-seq data from GM12878

The human EBV-transformed lymphoblastoid cell line (LCL) ATAC-seq data were downloaded from NCBI GEO database (GSE47753). The sequence reads from three replicates of 50k cell sample (GSM1155957, GSM1155958 and GSM1155959) were pooled and used for the downstream analysis. The 86,004 peaks called by MACS2(v2.1.1)^54,55^ were used for the downstream analysis.

## Acknowledgement

These studies were supported by funding from NHLBI (P01HL160476), Department of Defense (W81XWH2110606) and Minnesota Regenerative Medicine. We acknowledge the Minnesota Supercomputing Institute for providing computational resources.

